# Remodeled cephalic arch geometry promotes disturbed flow at pre-maturation flow rates in hemodialysis patients

**DOI:** 10.64898/2026.07.23.740371

**Authors:** Dylan Cook, Sanjeev Dhara, Maren Klineberg, Nhung Nguyen, Luka Pocivavsek, Bingqing Xie, Anindita Basu, Mary Hammes

**Author notes:** Corresponding Authors: Anindita Basu, Mary Hammes.

## Abstract

In the United States, roughly 550,000 people receive routine hemodialysis for end-stage renal disease. This treatment requires an arteriovenous access, most commonly a brachiocephalic fistula. However, these accesses often fail due to stenosis in the cephalic arch (CA), a common complication whose underlying causes remain unclear (Bennet *et al.*, 2015). To characterize the hemodynamic environment in the CA, we employed patient-specific millifluidic models that were perfused with blood-mimicking fluid containing fluorescently labelled beads to visualize flow behaviors. We perfused our models across physiologic (28-45 mL/min) and elevated (60-423 mL/min) flow rates and quantified wall shear stress (WSS) and streamline angle as a measure of flow disturbance. Our findings show that the bulk curvature and remodeled wall topography each create regions of persistently low WSS, consistent with prior clinical observations (Hammes *et al.*, 2016). Moreover, remodeled wall topography promotes disturbed flow at elevated flow rates, a hemodynamic profile associated with various vascular pathologies (Chiu & Chien, 2011). Independently performed computational fluid dynamics (CFD) modeling complements these results, showing that remodeled wall topography promotes vortex formation at elevated flow rates, as assessed by Q-criterion. Collectively, our experimental and computational results provide strong evidence for geometry-driven disturbed flow in the CA at elevated flow rates. Notably, we observed disturbed flow at flow rates as low as 81 mL/min, far below the 600 mL/min required for hemodialysis. Disturbed flow thus offers a plausible mechanism that relates access flow rates to the vascular pathologies that precede access failure.

**Significance Statement:** Hemodialysis requires an arteriovenous fistula (AVF) that must remodel and mature to withstand chronically elevated blood flow rates. However, how remodeled vessel geometry interplays with elevated flow to shape local hemodynamics remains poorly understood. Here, we used millifluidic models of the cephalic arch (CA) to show that vessel geometry and elevated flow rates promote regions of low wall shear stress and disturbed flow. Notably, geometry-driven disturbed flow is observed at flow rates above physiologic levels but well below 600 mL/min, the flow rate necessary for adequate dialysis. Because disturbed flow is injurious to the endothelium, our findings are the first to show that vascular damage begins before fistula maturation.

## Introduction

Roughly 550,000 people in the United States currently receive routine hemodialysis to treat end-stage renal disease (ESRD), a condition characterized by loss of at least 85% of kidney function (1). Hemodialysis requires a vascular access to both remove and return blood after filtering. Typically, this vascular access is an arteriovenous fistula (AVF) created by surgically connecting an artery and vein. The predominate AVF configuration is the brachiocephalic fistula (BCF) formed by connecting the brachial artery to the cephalic vein (2). However, BCFs have frequent clinical complications, with 30% of accesses failing after one year (3, 4). When an access fails, surgical or radiologic interventions are required to fix occlusions and/or create a new vascular access elsewhere in the body. Vascular access failure in BCFs is relatively common, leading to a median patency duration of about 3.5 years (5). Notably, between 39-77% of all patients with BCFs develop cephalic arch stenosis (CAS), which is the leading cause of access failure for these fistulas (3, 6–8). It is therefore crucial to understand the mechanisms underlying CAS.

Disturbed blood flow offers a plausible mechanism for CAS (9). Here, we define disturbed flow as deviations from laminar behavior, including eddy formation, flow separation from the vessel wall, and/or multidirectional flow (10). Under physiologic conditions, laminar blood flow exerts wall shear stress (WSS) against vessel walls that maintains endothelial morphology and homeostasis. By contrast, disturbed flow exerts low, oscillatory WSS, leading to endothelial reprogramming that promotes inflammation, apoptosis, and vascular remodeling (11). Consistent with these effects, disturbed flow has been linked to host of vascular pathologies. For example, an *ex vivo* model of a porcine saphenous vein showed that disturbed flow predisposed neointimal hyperplasia, a common precursor to venous stenosis (12–14). In humans, atherosclerotic plaques appear in reproducible regions along the vascular tree (e.g., aortic arch, valves, bifurcations) where disturbed flow is common (15–17). Collectively, this evidence demonstrates that disturbed flow precedes vascular pathologies across systems.

BCF placement and hemodialysis dramatically alter the hemodynamic environment of the CA. Before BCF creation, CA blood flow averages 28±14 mL/min (mean±SD) and the vessel diameter is 2-3 mm (18). Immediately upon BCF creation, blood flow through the CA increases significantly and the CA remodels to tolerate elevated blood flow rates, considerably modifying vessel geometry in the process (9, 19–21). In mature accesses, CA blood flow exceeds 600 mL/min and vessel diameter increases to 6-12 mm (22). Much previous work has sought to understand how these elevated blood flow rates and remodeled CA geometry influence local hemodynamics and patient outcomes. Early clinical observations associated high flow rates with CAS (23), while 2D computational models predicted geometry-driven disturbed flow and pockets of low WSS (24). Doppler ultrasound studies likewise suggested abnormally low WSS and high Reynolds number (*Re*) flow, though disturbed flow could not be directly confirmed (25, 26). Longitudinal studies further suggest that remodeling may transiently restore WSS, consistent with clinical and 3D computational evidence identifying the inner wall of the CA bend as a recurrent site of CAS and low WSS (27–30). Taken together, these findings indicate that abnormally high blood flow drives CA remodeling, giving rise to pockets of low WSS that predispose CAS.

These studies fall short of addressing fundamental questions regarding hemodynamics in the CA. Computational studies and Doppler ultrasound measurements suggest that disturbed, high-*Re* flow occurs, but they are unable to conclusively depict it. Moreover, it is unknown if the large-scale geometry of the CA (i.e., its diameter and curvature) determines its local hemodynamics or if small-scale vessel wall topography likewise has an effect. Understanding the effects of remodeled topographic features will reveal the degree to which patient-specific vessel geometry determines its hemodynamic environment. To address these questions, we previously developed patient-specific, millifluidic models of the CA to visualize local hemodynamics and calculate WSS, but these models were constrained to low flow rates (20 mL/min) (31). Here, we introduce an improved platform that allows us to observe hemodynamics from physiologic (28-45 mL/min) to elevated (60-423 mL/min) flow rates (**Fig. 1A**). We report varying degrees of disturbed flow across all patient models in both the bend of the CA and within small-scale topographic features at elevated flow rates. To complement our experimental evidence, we conducted computational fluid dynamic (CFD) to further examine the effects of vessel geometry on flow behavior. Overall, these findings offer conclusive evidence that remodeled CA geometry promotes disturbed flow and low WSS at elevated flow rates, possibly contributing to CAS and vascular access failure.

**Figure 1:**
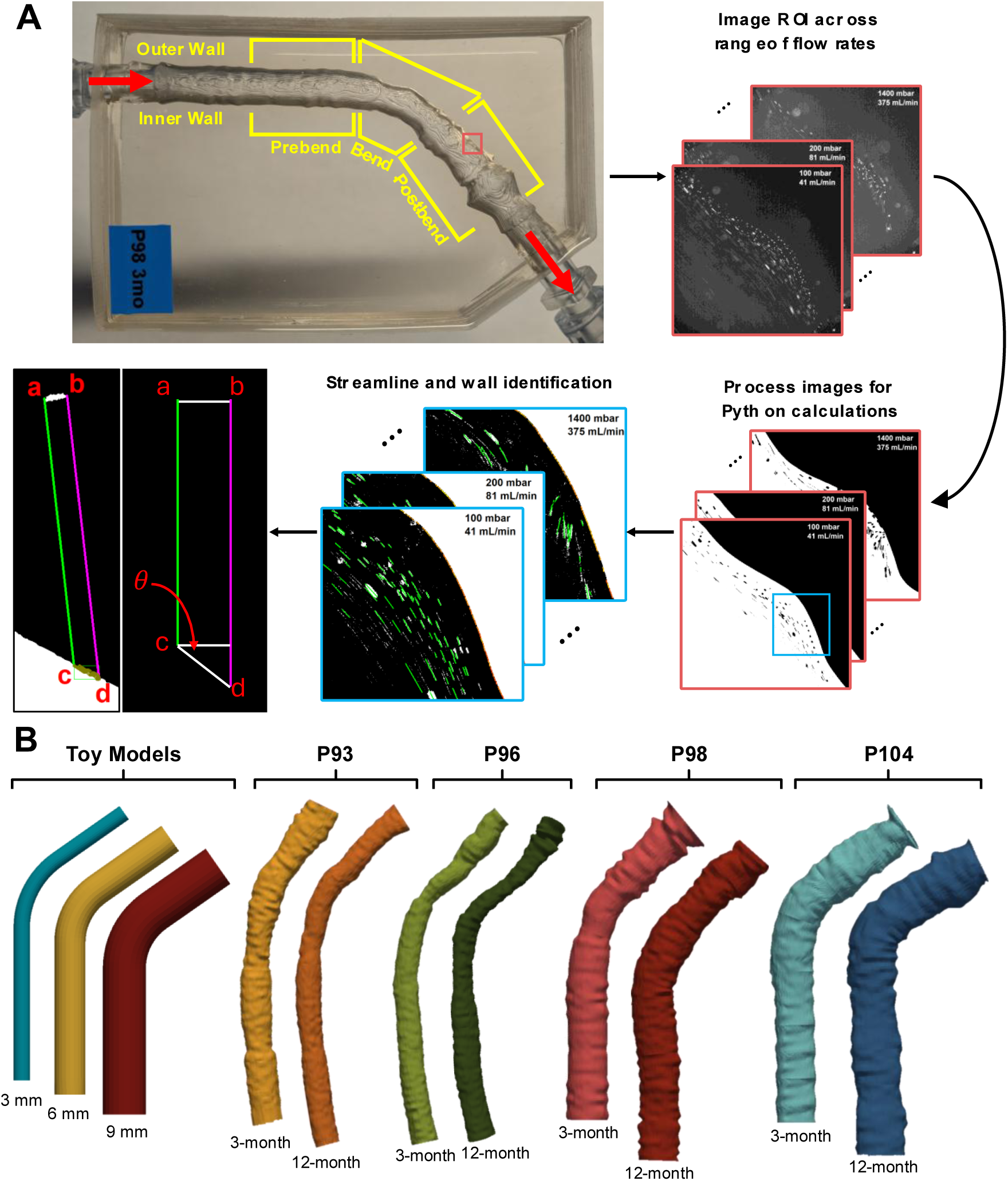
A strategy for characterizing local hemodynamics in millifluidic models of the cephalic arch. **(A)** Experimental overview for image acquisition, data processing, and calculations. Data is captured in millifluidic models perfused over a range of flow rates. Data are processed in ImageJ (NIH) and passed through a Python script to calculated wall shear stress and streamline angle relative to the wall. **(B)** CAD renderings of the toy and patient models used in this study.

## Results

### CA curvature promotes disturbed flow in large vessels at elevated flow rates

There are four primary features that may influence flow behavior in the hemodialysis patient CA: 1) elevated flow rates, 2) vessel curvature, 3) increased diameter, and 4) remodeled, uneven wall topography. Because wall topography varies dramatically between patients, we first isolated the effects of flow rate, vessel curvature, and vessel diameter using three toy models with topographically smooth walls and a 125° bend angle, representative of an average CA. The three diameters tested correspond to physiologic (3 mm) and enlarged (6 and 9 mm) CAs observed in patients with mature BCFs (**Fig. 1B**). Models were imaged across physiologic and elevated flow rates. We define elevated flow as >56 mL/min, which represents two standard deviations above the mean CA flow rate in individuals without BCFs. We adopted this cutoff to ensure that any effects attributable to flow rate occur only at rates firmly above physiologic norms. We hypothesized that the larger-diameter toy models would exhibit more disturbed flow, characterized by high streamline angle, and pockets of low WSS at elevated flow rates.

To determine how WSS varies with flow rate, we averaged the mean WSS values across regions of interest (ROIs) at each flow rate, producing a value we term global average WSS. As expected, global average WSS scaled linearly with flow rate and inversely with vessel diameter, with the 3- and 9-mm toy models exhibiting the highest and lowest values, respectively (**Fig. 2A**). However, WSS was distributed unevenly along the vessel in each model (**Fig. 2B-C**). To identify systematically low WSS regions, we selected ROIs whose mean WSS fell in the bottom decile of all (240-294 measurements) in ≥50% of tested flow rates (12-15 flow rates) (**Table 1**). In the 3- and 9-mm toy models, such ROIs consistently occurred along the inner wall downstream of the bend (**Table 1**). By contrast, in the 6-mm model, a systematically low WSS ROI was observed along the outer wall of the bend (**Table 1, Fig. S1B**). This is possibly due to higher pressure, and thus lower velocity, of the fluid as it rounds the bend, in accordance with Bernoulli’s theorem.

**Figure 2.**
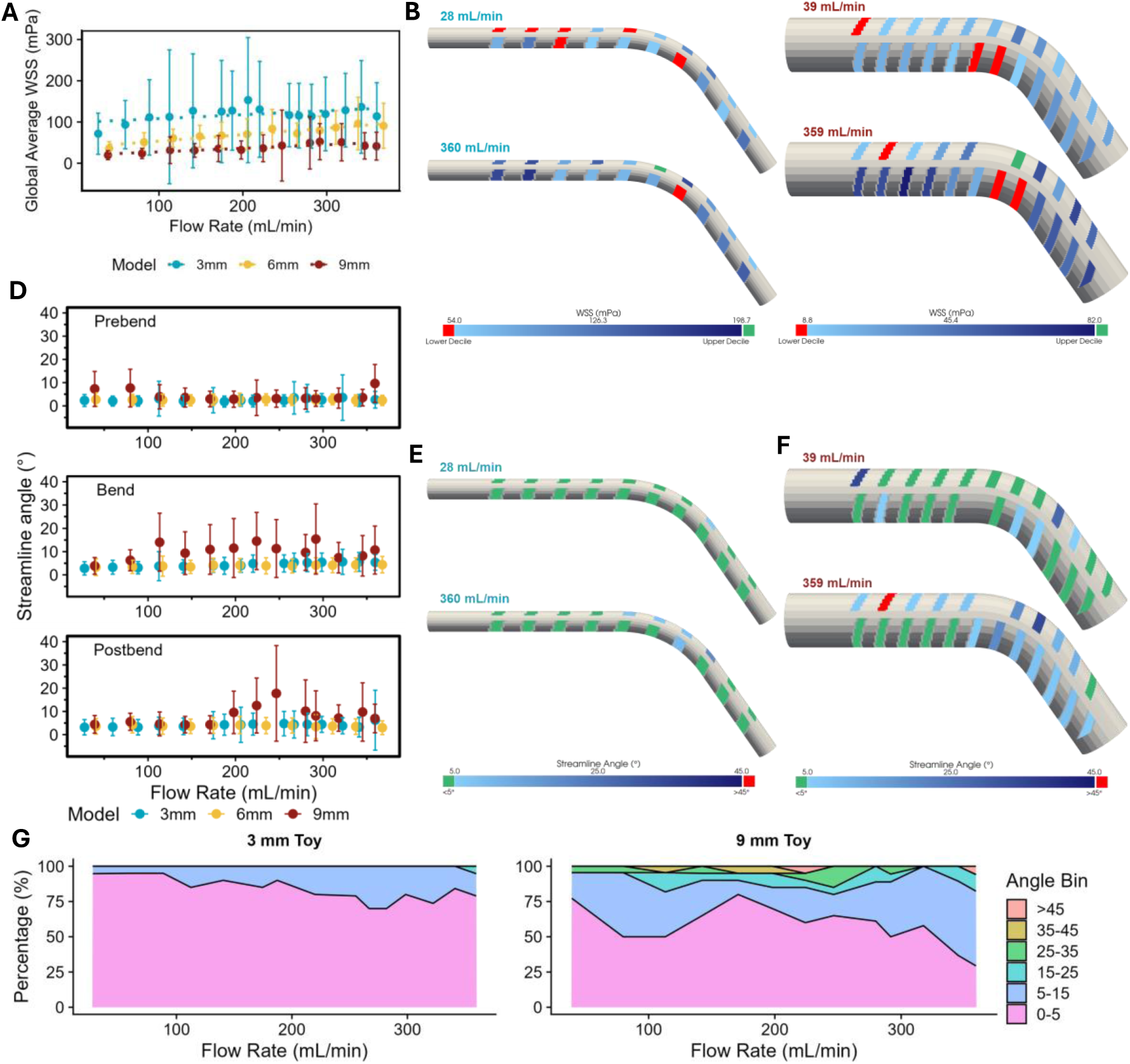
Disturbed flow develops around CA curvature at high flow rates. **(A)** Global average wall shear stress (WSS) scales with flow rate in three idealized toy models. **(B-C)** 3 and 9 mm toy models colored with WSS magnitudes measured at indicated flow rates. **(D)** Domain-averaged streamline angle versus flow rate in three toy models (3, 6, and 9 mm diameter). Models are segmented into three domains (prebend, bend, postbend). **(E-F)** 3 and 9 mm toy models colored with measured streamline angle values. **(G)** Proportion of ROIs that fall into designated angle ranges.

**Table 1.**
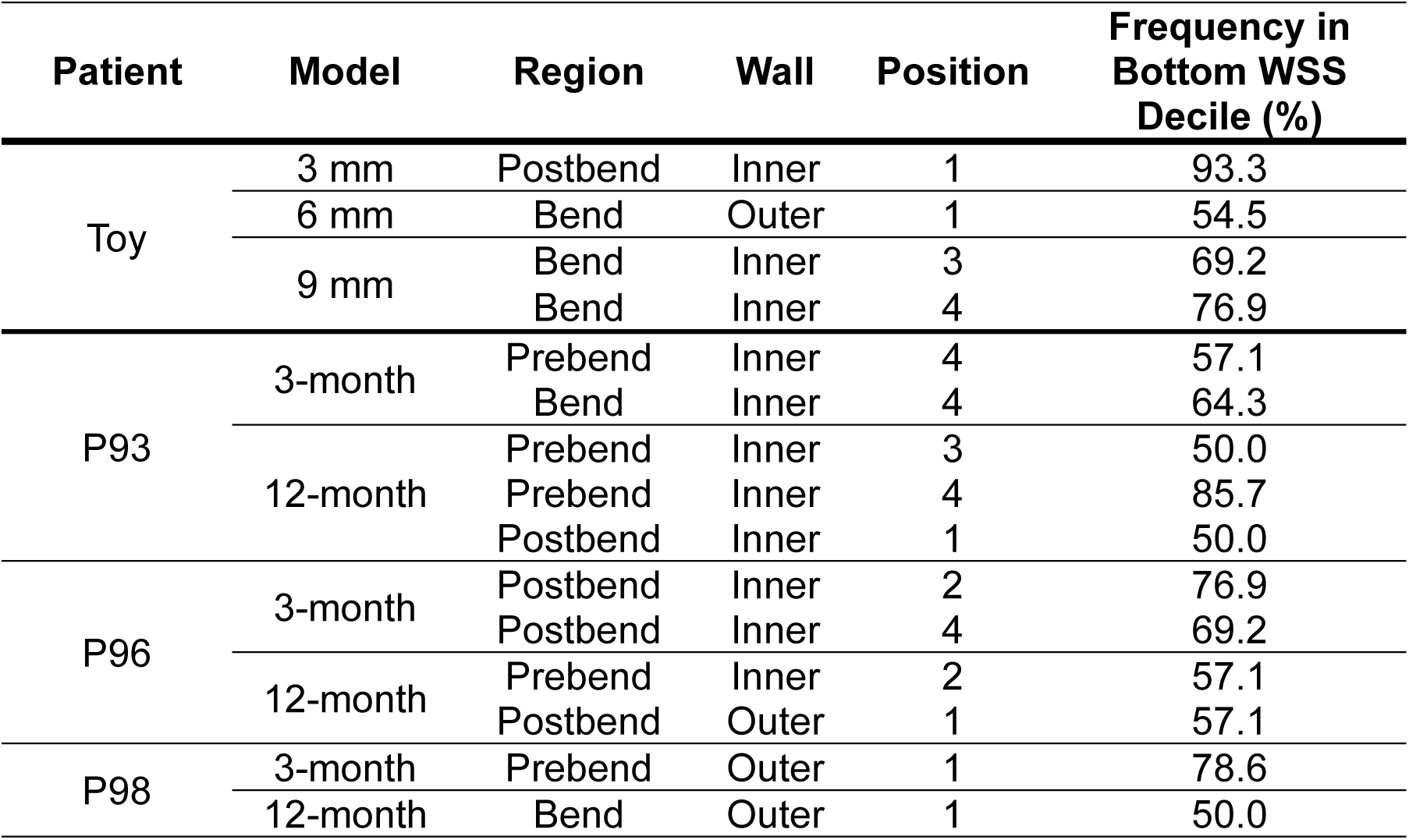
Systematically low WSS zones in toy and patient models.

| Patient | Model | Region | Wall | Position | Frequency in Bottom WSS Decile (%) |
| --- | --- | --- | --- | --- | --- |
| Toy | 3 mm | Postbend | Inner | 1 | 93.3 |
|  | 6 mm | Bend | Outer | 1 | 54.5 |
|  | 9 mm | Bend | Inner | 3 | 69.2 |
|  |  | Bend | Inner | 4 | 76.9 |
| P93 | 3-month | Prebend | Inner | 4 | 57.1 |
|  |  | Bend | Inner | 4 | 64.3 |
|  | 12-month | Prebend | Inner | 3 | 50.0 |
|  |  | Prebend | Inner | 4 | 85.7 |
|  |  | Postbend | Inner | 1 | 50.0 |
| P96 | 3-month | Postbend | Inner | 2 | 76.9 |
|  |  | Postbend | Inner | 4 | 69.2 |
|  | 12-month | Prebend | Inner | 2 | 57.1 |
|  |  | Postbend | Outer | 1 | 57.1 |
| P98 | 3-month | Prebend | Outer | 1 | 78.6 |
|  | 12-month | Bend | Outer | 1 | 50.0 |

To quantify flow disturbance, we calculated the mean angle between streamlines and the vessel wall—a measure we term “streamline angle”—to assess how far flow deviates from laminar (i.e. streamline angle = ∼0°) (**Fig. 1A**). Streamline angles of 0-5° were classified as low-angle flow (minimally disturbed) and angles >5° as high-angle flow, indicative of flow disturbance. We mark this threshold to enable comparisons rather than to distinguish an exact physiologic transition. First, mean streamline angles were averaged across ROIs within the prebend, bend, and postbend domains to observe domain-specific effects across flow rates (**Fig. 2D**). In all toy models, the prebend domain remained largely laminar across all flow rates. This trend extended into the bend and postbend domains for the 3- and 6-mm toy models. By contrast, the 9-mm toy model displayed increased streamline angle in the bend and postbend domains as flow rate increased. As with WSS, streamline angles varied along the length of the model (**Fig. 2E-F**). To capture global trends, ROIs were binned by mean streamline angle (**Fig. 2G**). In this schema, a high proportion of ROIs experiencing low-angle flow indicates a low degree of disturbed flow, and vice versa. Broadly, the proportion of ROIs experiencing low-angle flow was higher in the 3-mm model than the 9-mm model across all flow rates. In both models, the proportion of ROIs with low-angle flow decreased as flow rate increased, suggesting progressively greater flow disturbance.

### Remodeled vessel topography promotes disturbed flow in patient models

Because the toy models had smooth vessel walls, we next examined whether remodeled, patient-specific wall topography altered local hemodynamics. We compared the P96 patient models at 3- and 12-month timepoints (mean diameters = 6.6 mm; bend angles = 133° and 132°, respectively) with the geometrically similar 6-mm toy model (bend angle = 125°), attributing observed differences in flow behavior to surface topography. As in the toy models, global average WSS in the P96 models scaled linearly with flow rate (**Fig. S1A**). While the P96 models likewise harbored systematically low WSS zones, these zones primarily occurred in the prebend and postbend domains of the model rather than the bend (**Table 1**). In contrast to the 6-mm toy model, which globally maintained low-angle flow across flow rates, the P96 models displayed increased streamline angles in all domains, with the highest values observed in the bend and postbend (**Fig. 3A-D**). In the 6-mm toy model, >75% of ROIs retained low-angle flow across all flow rates; the P96 patient models were susceptible to high-angle flow at mildly elevated flow rates (>56 mL/min) and predominated by it above ∼150 mL/min (**Fig. 3E**). This trend was repeated among the other patient models (**Fig. S2-4**). Together, these findings demonstrate that remodeled vessel topography substantially increases the amount of disturbed flow in the CA at elevated flow rates.

**Figure 3.**
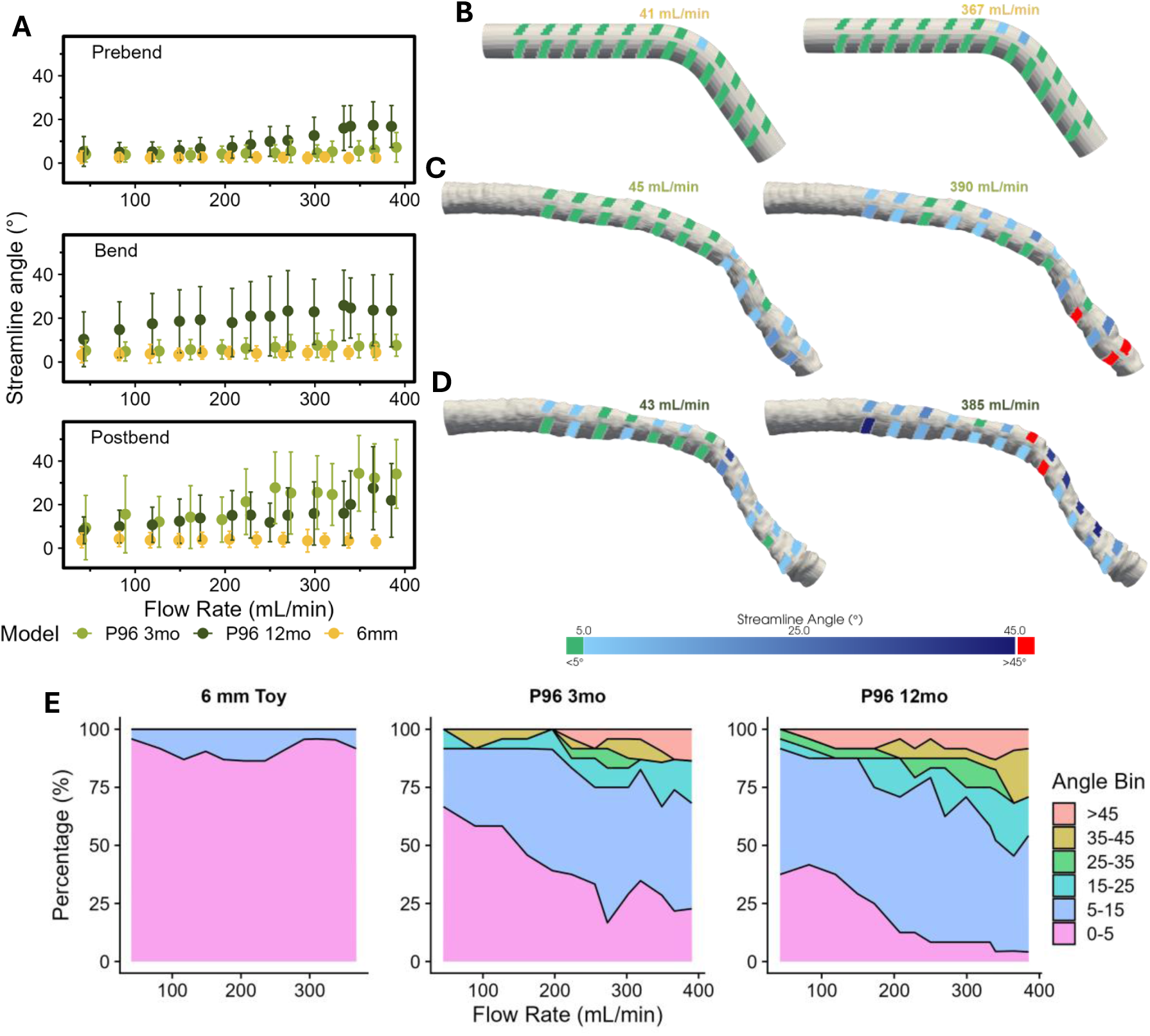
Remodeled vessel topography promotes disturbed flow in patient models. **(A)** Domain-averaged streamline angle versus flow rate comparing 6 mm toy and P96 patient (3- and 12-month timepoints) models. **(B-D)** Streamline angle magnitudes colored onto 6 mm toy **(B)**, P96 3-month **(C)**, and P96 12-month **(D)** models. **(E)** Proportion of ROIs that fall into designated angle ranges.

### Large- and small-scale vessel features promote disturbed flow

We observed recurrent non-laminar flow patterns associated with specific vessel features across the toy and patient models (**Fig. 4**). In the 9-mm toy model, the inner wall of the bend harbored a consistently low WSS zone. This site was laminar at a physiologic flow rate (39 mL/min) but prone to eddy formation at moderately elevated values (224 mL/min) (**Fig. 4A**). We also observed similar low WSS at elevated flow in patient models with comparably low-radius bends, including P93 3-month, P98 12-month, and P104 12-month (**Figs. S2D, S3G, S4G**). Additionally, gradual postbend diameter expansion was associated with streamline angles increasing linearly with flow rate, yielding flow that appeared orthogonal to the vessel wall (**Fig. 4C**).

**Figure 4.**
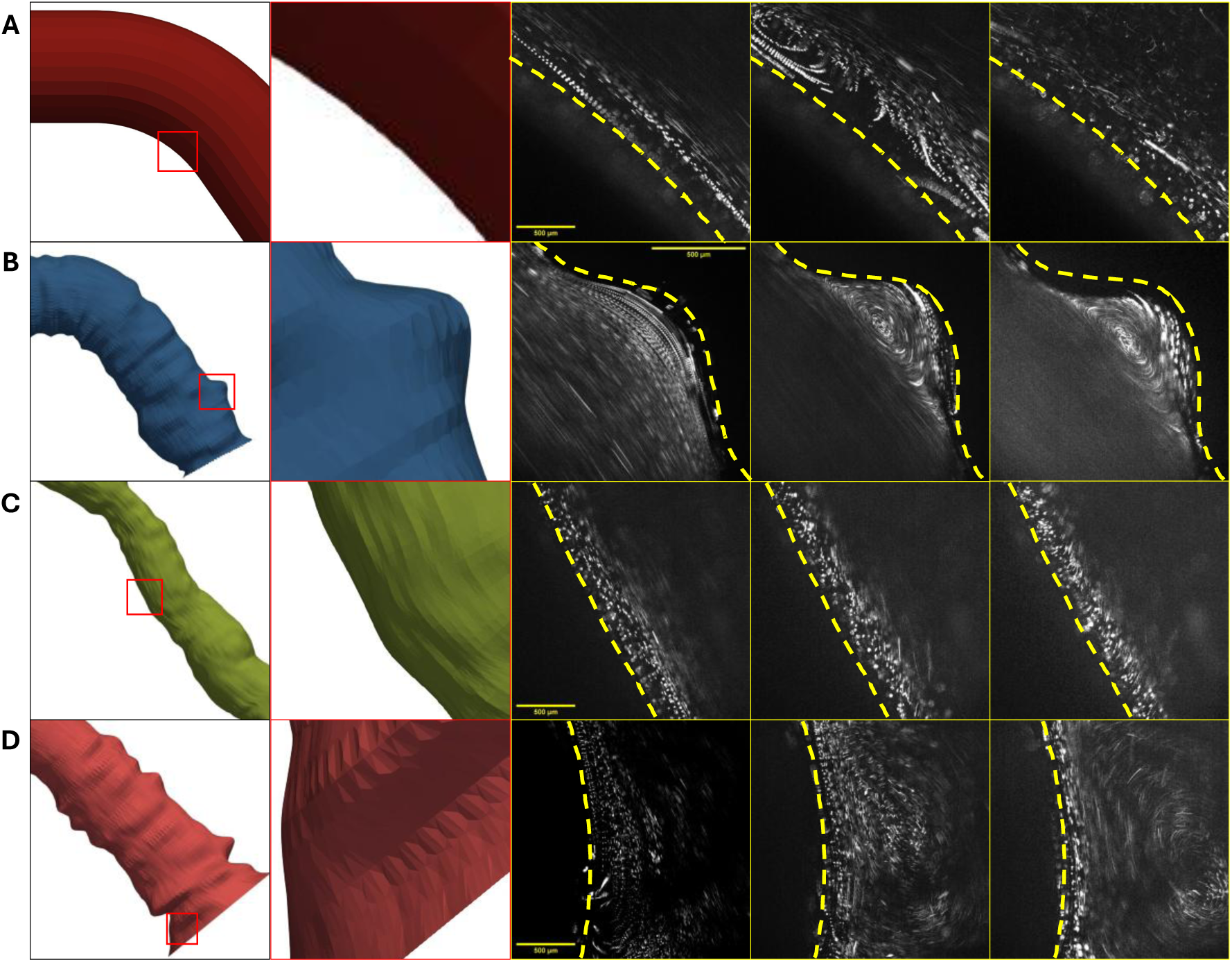
Large- and small-scale geometric features influence disturbed flow development. **(A)** At sufficiently large diameters, innate cephalic arch curvature causes an eddy to develop at high flow rates. **(B)** As fluid passes over a topographic depression, a low-velocity, recirculating eddy forms within the bulge while the adjacent flow remains laminar. **(C)** Along gradual increases in vessel diameter, streamline angle gradually increases as flow rate increases. **(D)** Instances where vessel diameter narrows and expands over short distances results in eddies at the site of vessel expansion.

Local variations in wall topography also corresponded to isolated regions of disturbed flow. Pronounced bulges in the vessel wall (e.g. observed in P104 12-month) remained laminar at physiologic flow rates but were susceptible to eddy formation at mildly elevated ones (**Fig. 4B, Movie S1**). Similar bulges were present in P96 12-month and P98 3-month patient models. Finally, some patient models displayed short-segment diameter narrowing, reminiscent of an early stenosis. At mildly elevated flow (∼81 mL/min), the post-narrowing expansion harbored an eddy (**Fig. 4D, Movie S2**). Overall, these observations demonstrate how large- and small-scale features in remodeled vessels promote flow disturbance in patient-specific models.

### Remodeled topography increases disturbed flow along vessel walls

We performed computational fluid dynamics (CFD) modeling in XFlow to complement our experimental findings and characterize disturbed flow in regions missed by our imaging protocol. We compared the 9-mm toy model to the P98 3- and 12-month timepoints because they are similar in both average diameter (8.2 and 10.3 mm, respectively) and bend angle (130° and 115°, respectively). We simulated flow at physiologic (40 mL/min), moderately elevated (200 mL/min), and highly elevated (360 mL/min) flow rates. Additionally, for the P98 3- and 12-month models, we performed simulations at systolic flow rates (1586 and 887 mL/min, respectively) measured by Doppler ultrasound (30). From each simulation, we extracted the Q-criterion, which quantifies regions where the rotational components of the velocity gradient tensor predominate over the strain components (**Fig. 5**) (32). As such, regions of positive Q-Criterion highlight regions predisposed to vortex formation. We hypothesized that the patient models would display higher values of Q-criterion, indicative of greater flow disturbance, due to their uneven wall topographies (33). Overall, the 9-mm toy model shows minimal areas of positive Q values, indicating a low degree of flow disturbance (**Fig. 5**). By contrast, both P98 models show regions of alternating positive and negative values of Q-criterion (**Fig. 5**), notably localized to the vessel walls of the prebend domain. In addition, we note larger regions of positive Q values in the bend and post-bend regions due to vessel curvature. These regions first appear at moderately elevated flow rates (∼200 mL/min) and are exacerbated at higher (360 mL/min) and at patient-specific values. The velocity profiles at the aforementioned conditions are shown in **Fig. S5**.

**Figure 5:**
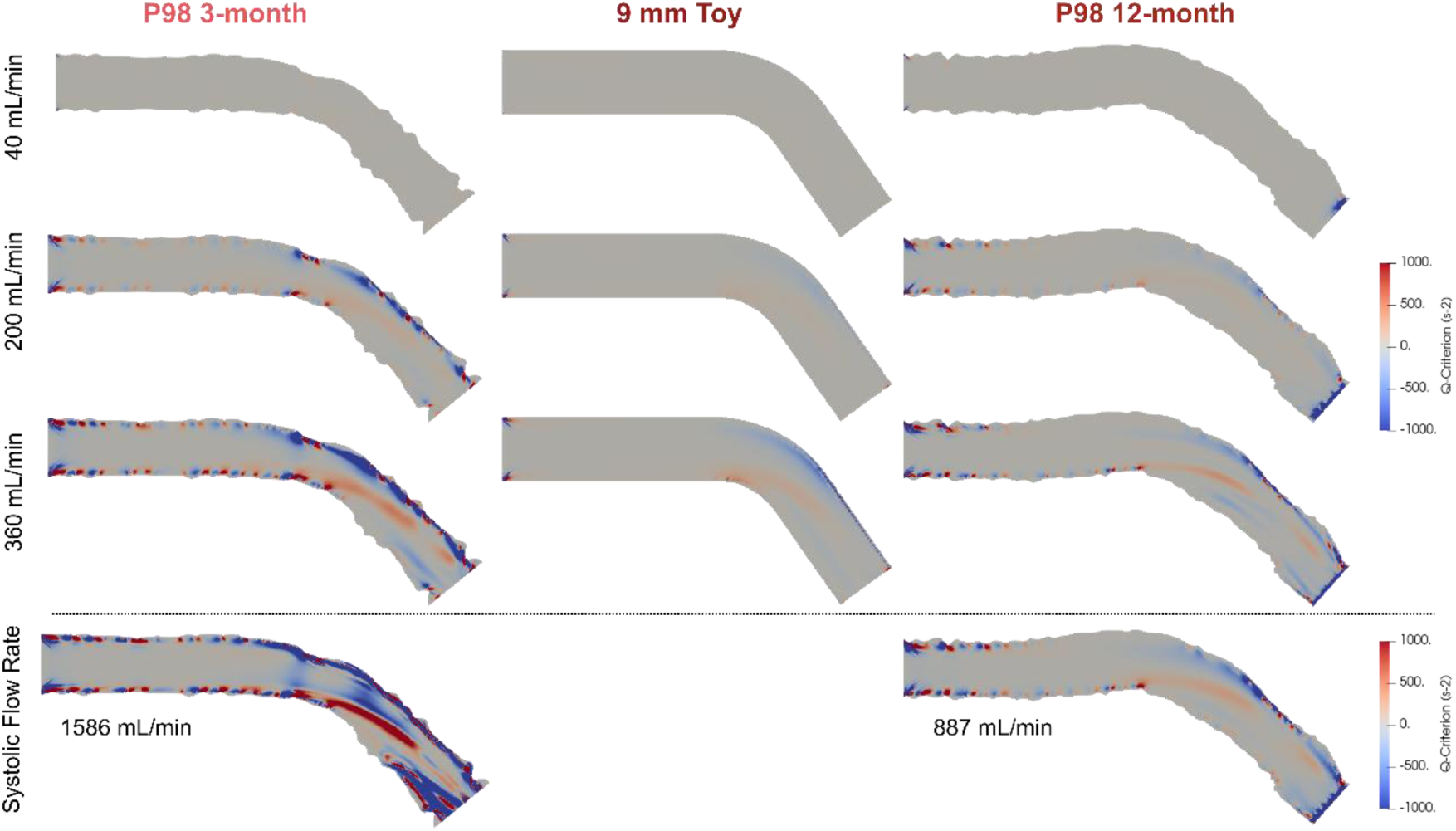
Remodeled topography increases flow disturbance along vessel walls. Distribution of Q-criterion in P98 3-month, 9 mm toy, and P98 12-month models. Computational fluid dynamics simulations were performed at physiologic (40 mL/min), moderately elevated (200 mL/min), and highly elevated (360 mL/min) flow rates. For the patient models, additional simulations were performed at observed systolic flow rates.

To observe the circumferential distribution of Q-criterion at 360 mL/min, we plotted values at discrete slices along the lengths of the models (**Fig. S6**). This perspective shows that the 9 mm toy model developed disturbed flow in the center of the model in and after the bend (60-85% through the model), which agrees with our physical models (**Figs. 2C, 4A**). In contrast to the 9 mm model, the P98 3-month model shows a markedly higher degree of disturbed flow, again concentrated in and after the bend. The P98 12-month model likewise shows disturbed flow in the same regions, albeit to a lesser degree than its 3-month counterpart (**Fig. S6**).

## Discussion

Previous studies have established that elevated blood flow rates correlate with the development of CAS in hemodialysis patients, but how elevated flow interplays with remodeled vessel geometry to shape local hemodynamics remains poorly understood. In this study, we implemented patient-specific millifluidic models of the CA to characterize the spatial distribution of wall shear stress (WSS) and disturbed flow over a range of physiologic and elevated flow rates. By quantifying WSS and streamline angle across flow rates and remodeled geometries, our platform demonstrates how large-scale vessel geometry and local, small-scale topographic features each interact with elevated flow rates to yield reproducible pockets of disturbed flow and low WSS in patient models.

Across toy and patient models, we observed reproducible zones of low WSS in all major anatomical domains. Many of these zones remained in the bottom decile of WSS measurements across all flow rates tested, even though global average WSS increased with flow. Because chronically low WSS in the CA is associated with stenosis (29), our data provide evidence for geometry-driven susceptibility. Strikingly, the single ROI most frequently (93.3%) found in the bottom decile of WSS measurements resided in the early postbend of the 3 mm model, which most closely replicates natives CA geometry. Therefore, pockets of low WSS are likely an inherent consequence of CA geometry that are exacerbated post-BCF creation and maturation.

By calculating streamline angle, we developed a method of quantifying flow disturbance without assuming topographically smooth vessels, as is required for Reynolds number (*Re*) calculations. Validation in toy models showed low-angle flow in the prebend domain, which effectively mimics a smooth, straight pipe. In sufficiently large vessels (9 mm), the inner wall of the bend was prone to disturbed flow at moderately elevated flow rates. Thus, in the absence of topographic irregularities, CA curvature alone is sufficient to promote disturbed flow at elevated flow rates. In our patient models, streamline angle measurements further identified local geometric features, such as bulges and short-segment narrowing, that harbored disturbed flow at mildly elevated flow rates (81 mL/min). CFD simulations corroborated our physical models. High Q- criterion values along remodeled vessel walls reiterated our observations that local topographic features promote disturbed flow. Interestingly, we observed an improvement of flow disturbance between the P98 3- and 12-month timepoints due to smoother vessel walls in the latter, possibly indicating adaptive remodeling over time (27). Overall, we conclude that remodeled vessel geometry cooperates with elevated flow rates to promote disturbed hemodynamics, complementing prior Doppler ultrasound evidence of high-*Re*, turbulent flow in the CA of hemodialysis patients (25, 26).

### Limitations

In this study, we established millifluidic models as a robust platform for quantifying WSS and streamline angle and for qualitatively assessing small-scale flow behavior. These models offer a significant improvement over previous versions, but are still limited in several key aspects. The PDMS models do not match the viscoelastic properties of human blood vessels and therefore cannot capture vessel deformations over a cardiac cycle. Nevertheless, patient models agree well with their *in vivo* counterparts, as assessed by IVUS (31). Similarly, all measurements were all taken under steady-state flow and do not capture pulsatility, which typically exacerbates disturbed flow during cyclic acceleration and deceleration. Technical constraints also limited maximum flow rates to 350-400 mL/min, which fall below the 600 mL/min necessary to support adequate hemodialysis (22). We believe that these technical limitations cause our system to underestimate the hemodynamic environment that may be found *in vivo*. Finally, the CFD models are limited by assuming rigid walls and Newtonian properties for blood.

## Conclusion

This study offers clinically important insights into how remodeled CA geometry interacts with elevated flow rates to shape local hemodynamics. Using patient-specific millifluidic models and complementary CFD simulations, we demonstrate that CA curvature and remodeled vessel topography generate persistent regions of low wall shear stress (WSS) and disturbed flow— hemodynamic conditions associated with vascular remodeling, inflammation, and coagulation. Although the elevated flow rates required for hemodialysis are necessary for proper treatment, our findings show that elevated flow in the context of remodeled vessels promotes regions of low WSS and disturbed flow. These hemodynamic features are associated with endothelial dysfunction and vascular pathologies, offering a plausible mechanistic link between access flow and access failure. While evidence suggesting adaptive remodeling was observed in one patient between their 3- and 12-month timepoints, this adaptation was insufficient to restore laminar, undisturbed flow. Most strikingly, geometry-driven disturbed flow was observed in patient models at flow rates as low as 81 mL/min. Because these flow rates fall far below 600 mL/min, the flow rate required for adequate dialysis, disturbed flow and consequent damage to the endothelium likely occur before fistula maturation. Our findings are the first to demonstrate that vascular damage first occurs between fistula placement and maturation, shifting the timeline of pathogenesis up to before hemodialysis begins. In summary, high blood flow rates drive the development of pathologic flows in the CA in a geometry-specific manner, making blood flow rate a key consideration in assessing patient risk for vascular complications (34).

## Materials and Methods

### Device fabrication

The device fabrication protocol was adapted to create devices that could withstand elevated flow rates up to ∼400 mL/min (31). To study cephalic arch (CA) hemodynamics in patients, 3D models of the CA of four hemodialysis patients at two time points (3 and 12 months after AVF placement) were created from previously published IVUS and venogram data (**Fig. 1B**) (30). Models were imported into Trimble SketchUp and modified to add barbed tubing adapters and close geometries. In addition to the patient-specific models, three idealized “toy” models were created to study how vessel curvature and diameter alone affect local hemodynamics in the CA. To this end, we created the toy models with average diameters of 3 mm, 6 mm, and 9 mm, each with a bend angle of 125°.

Once the 3D models were created, they were exported as *.stl files and prepared for 3D printing in the Cura LulzBot Edition 3.6.25 software as GCode files (*.gcode). Models were printed on a Taz4 3D printer (B&H Photo, #LUKTPR0041NA) in polyvinyl alcohol filament (PVA; eSUN, #PVA300N05) at 100% model density and 10% support density. Following 3D printing, the surfaces of the models were smoothed by gently wetting them with crushed ice and wiping away partially dissolved PVA to mitigate layer artifacts without altering the geometric features of the models. Finally, a perimeter was traced around each model to 3D print a box mold.

Next, the 3D-printed models were cast in polydimethylsiloxane (PDMS; Dow Sylgard 184, #4019862). Each box mold was affixed to a single-well polystyrene plate (Sigma-Aldrich, #O0764) with a hot glue gun. PDMS was mixed in a mass ratio of 1:10 (crosslinker : base) inside a Thinky AR-100 conditioning mixer. Mixed PDMS was poured into the box mold to form a base layer, degassed in a vacuum desiccator for 30 min, and partially cured at 65°C for 1 hr. This degassing and partial curing process was repeated for three additional layers: PDMS was poured to (1) embed the 3D-printed CA model halfway, (2) just cover the model, and (3) fully encase the model. After the final layer, devices were fully cured at 65°C for 2 hr, after which the box mold and Petri dish were removed to reveal the PDMS-encased CA model.

To finish device fabrication, the internal 3D-printed CA model was dissolved and tubing was attached. A 3 mm biopsy punch was used to open both ends of the devices. The flat surfaces of the models were then covered with aluminum PCR foil (ThermoFisher, #AB0626) to protect them from residues during the dissolving step. PVA was dissolved from the models by submerging them in tap water and successively autoclaving the devices in a B4000-16 BioClave Research Autoclave (Benchmark Scientific) at 134°C and 30 psi until no trace of PVA remained. Next, the inlet and outlet tubing were prepared. Two 10 cm lengths of 1/4” ID tubing (McMaster-Carr, #5393K41) were fitted with a 1/4” barbed tube fitting (McMaster-Carr, #5117K46) on one end and a 1/4” × 1/8” ID barbed tube fitting (McMaster-Carr, #5117K61) on the other. PDMS was prepared as previously described and gently applied to the outer surface of the 1/4” barbed tube fitting. This end of the tubing was then inserted into each end of the device, where it locked into the barbed fitting element of the 3D design. Devices were then incubated at 65°C for 2 hr to cure the PDMS and secure the tubing in place.

### Device perfusion

Devices were perfused via pressure-driven flow supplied by an Elvesys OB1 MK3 control system attached to an air compressor (California Air Tools, #2010A). Applied pressure drove a blood-mimicking fluid (BMF, 6.3% dextran in distilled water [w/v], Sigma-Aldrich, #D4876), which closely approximates the viscosity and density of blood (3.5 mPa·s) from a fluid reservoir through the devices (31). For flow visualization, BMF was supplemented with 3.5 × 10^-6^ % (v/v) of 2.0 µm, fluorescently labelled polystyrene beads (Invitrogen, #F8827). BMF traveled through 1/16” ID tubing (McMaster-Carr, #6519T16), transitioned into 1/8” ID tubing (McMaster-Carr, #5238K718) via a barbed tube fitting (McMaster-Carr, #5117K52), and finally connected to the device inlet. A length of 1/8” ID tubing at ambient pressure was used as outlet tubing.

### Image Acquisition

Our image acquisition setup has been previously described (31). Devices were imaged on an Olympus IX83 with a Hamamatsu ORCA Flash4.0 camera and MetaMorph software (Molecular Devices). Unless otherwise noted, timelapse images were taken with 6.4X magnification (4X, NA = 0.16 objective with 1.6X mirror magnification), 40 fps, 100 ms exposure time, and GFP illumination (488 nm/510 nm). Each model was subdivided into prebend, bend, and postbend domains (**Fig. 1A**), which were further partitioned into smaller regions of interest (ROIs). Both the inner and outer walls of each ROI were imaged. Per ROI, 50 frames were captured across steady-state flow rates created by increasing the applied pressure from 100 to 1200-1600 mbar in 100 mbar increments. This range encompassed physiologic (28-45 mL/min) and elevated (60-423 mL/min) flow conditions. Flow rates at each applied pressure value were determined by measuring the collection time of 20 mL effluent.

### Image Processing

Raw image data was processed for qualitative visualization and wall shear stress (WSS) and streamline angle calculations. All image processing was performed with built-in functions in NIH ImageJ; all custom macros are publicly available. For qualitative presentation, 50-frame stacks from each ROI and flow condition were brightness- and contrast-enhanced before creating a maximum intensity Z-projection. For WSS and streamline angle calculations, vessel walls and streamlines were processed separately. Wall contours were manually traced and binarized from maximum-intensity projections, while streamlines were isolated via background subtraction, thresholding (Shanbhag), and particle analysis. The vein wall and streamline masks were then combined to generate binary image stacks for downstream analysis.

### WSS & Streamline Angle Calculations

Wall shear stress (WSS) was calculated using a previously reported Python script (31). In short, streamline lengths and wall distances were measured in OpenCV. We computed WSS using the formula 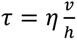, where *η* is viscosity, *v* is particle velocity, and ℎ is the streamline-wall distance.

Mean WSS was calculated per ROI, and values were averaged across ROIs at each flow rate to obtain a value we term global average WSS.

Streamline angle relative to the wall was calculated to quantify flow disturbance. Streamline endpoints (A and B) were used to determine streamline orientation, and lines perpendicular to the streamline were extended to intersect the wall contour (points C and D; **Fig. 1A**). The lengths of line segments *AB*, *AC*, and *BD* were used to compute the streamline angle, *θ*, relative to the wall using the formula: 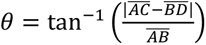. Mean streamline angles were calculated per ROI and further averaged with each domain to obtain a value we term domain-averaged streamline angle.

### Computational Fluid Dynamic Modeling

Design (*.stl) files of the 6 and 9 mm toy models and patient P98 3-month and 12-month models were used for computational fluid dynamic (CFD) modeling. Geometries were pre-processed in Simpleware ScanIP (Synopsys, Exeter, United Kingdom) to ensure watertight surfaces and apply a minimal recursive Gaussian smoothing filter to reduce any jagged surface triangles. The processed *.stl files were then exported to Blender 4.4 (Blender Foundation) to identify boundaries. A computational model was constructed in XFlow (Simulia, Dassault Systems). Blood was modelled as a Newtonian fluid with a dynamic viscosity of 3.5 mPa·s and density of 1060 kg/m^3^. To isolate the impact of the differing surface topography between the models, simulations were run under three uniform inlet volumetric flow rates: 40, 200, and 360 mL/min. The total simulation time was 1 second and the time step was 2.5 × 10^-6^ s. A uniform lattice discretization of 150 µm was applied. We utilized 20 CPU cores for each model and computing resources from The Center for Research Informatics at the University of Chicago Biological Sciences Division. The velocity field and Q-criterion were extracted from the simulations for further analysis.

## Author Contributions

D.C.: designed and performed research, analyzed data, wrote the paper.

S.D.: performed research, analyzed data, wrote the paper.

M.K.: designed and performed research, analyzed data.

N.N.: performed research, analyzed data.

L.P.: reviewed the paper.

B.X.: contributed analytical techniques.

A.B.: designed research, analyzed data, wrote the paper.

M.H.: designed research, wrote the paper.

## Acknowledgments

We extend our deepest gratitude to the patients whose IVUS data was used in this study, without whom this work would not be possible. We thank Linus Hansen and Alexander Katz for their help with data collection. We would also like to thank Frederick Naumann, Natalia Gonzales, and Sarah Sumner for their help in preparing the manuscript. We would also like to acknowledge The Center for Research Informatics, which is funded by the Biological Science Division, USA at the University of Chicago. This work was supported in part by the GI Research Foundation Young Investigator Award to N.N., which supported the time and effort of N.N. and S.D., and by the NSF under Award No. CMMI-2433223 to N.N., which supported computational resources and development of the fluid simulation workflow used in this work. Generative AI (GPT-4) was used to correct code and produce feedback for manuscript revisions.

---

**Figure S1:**
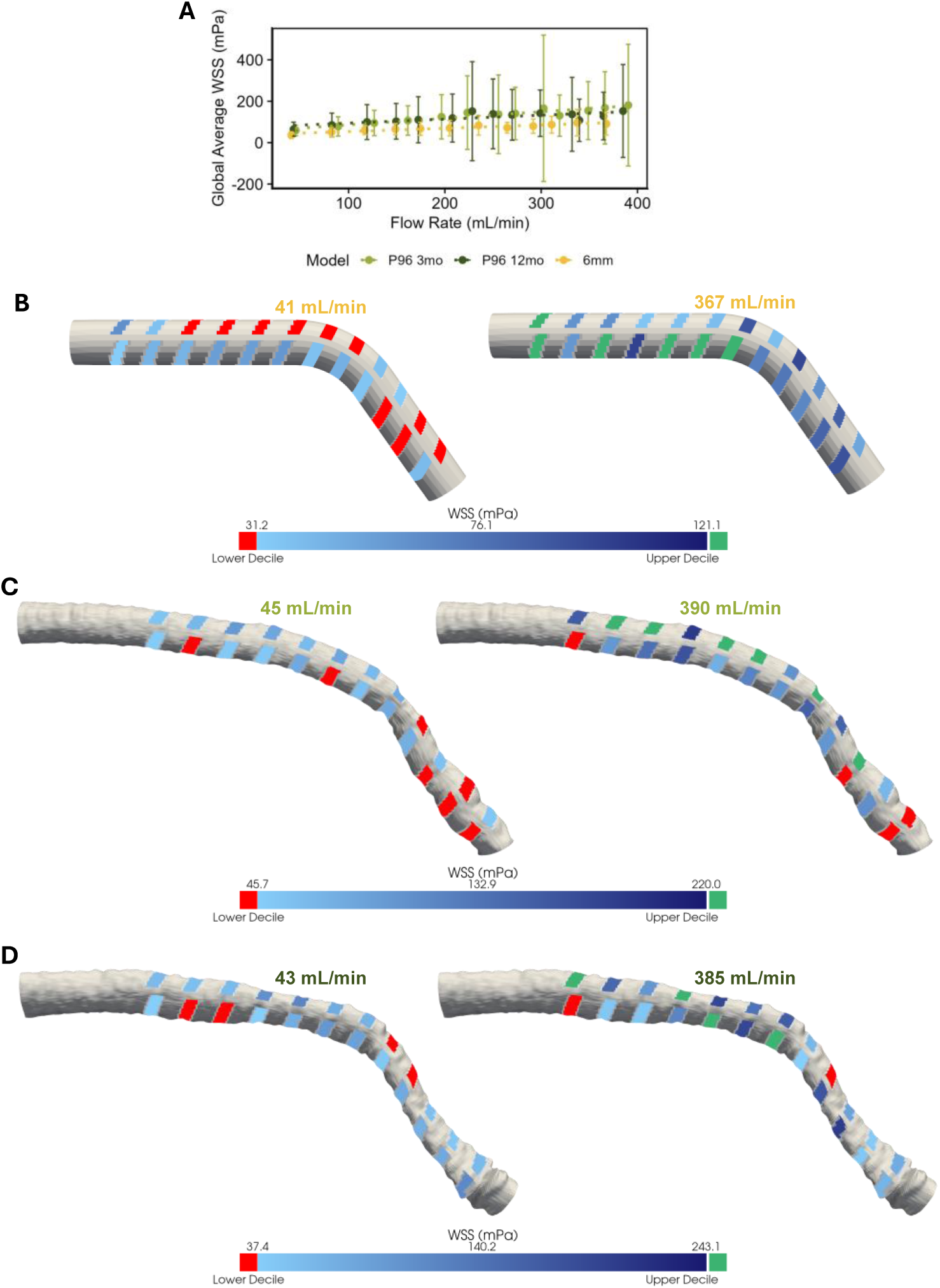
WSS profiles of 6 mm toy and P96 patient models. **(A)** Global average wall shear stress (WSS) scales with flow rate. **(B)** 6 mm toy, **(C)** P96 3-month, and **(D)** P96 12-month models colored with WSS magnitudes measured at indicated flow rates.

**Figure S2:**
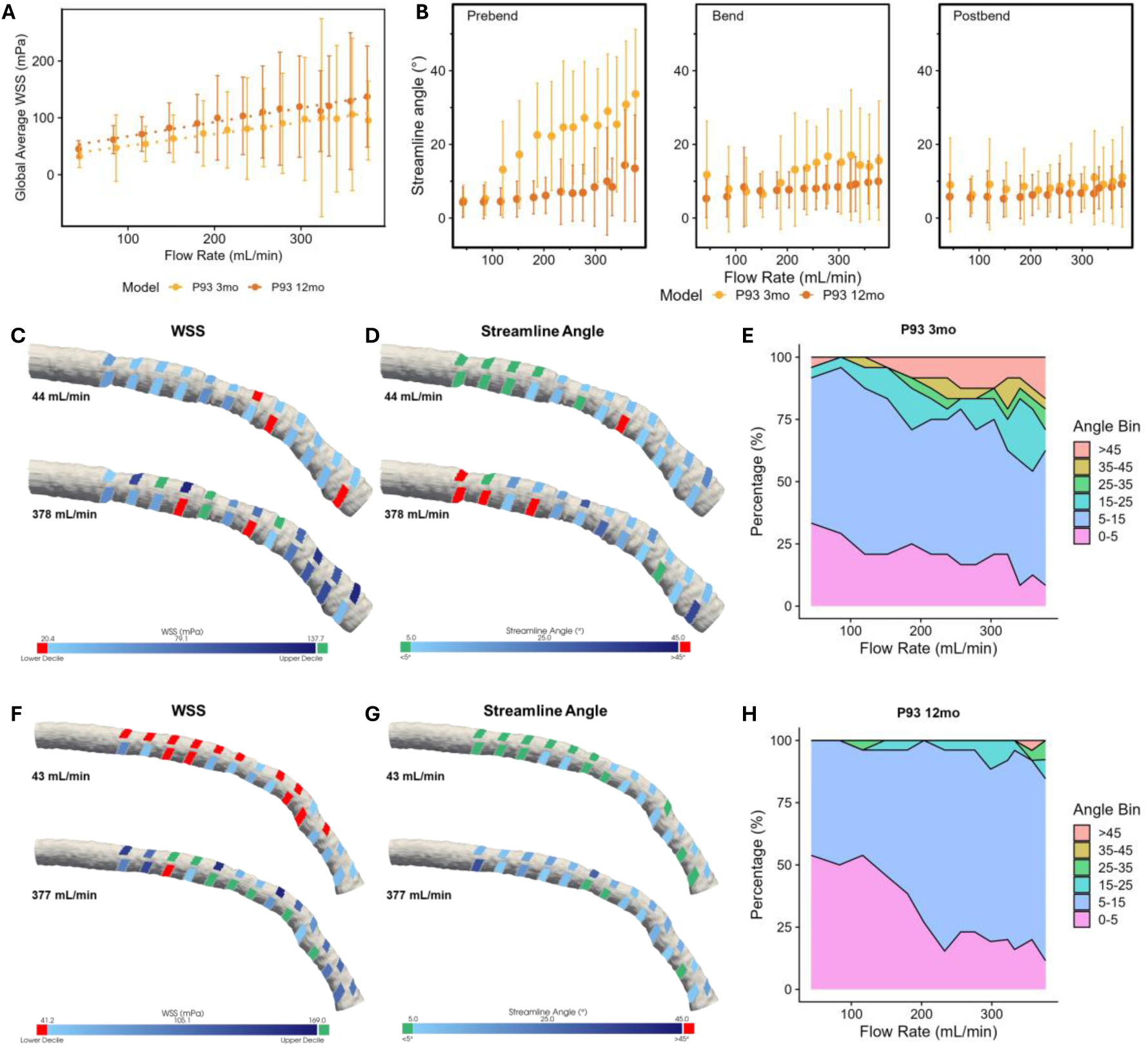
Hemodynamic profile of P93 patient models. **(A)** Global average wall shear stress (WSS) scales with flow rate. **(B)** Region-averaged streamline angle versus flow rate. Models are segmented into three regions (prebend, bend, postbend). **(C,F)** WSS values projected onto 3-month **(C)** and 12-month **(F)** models at physiologic and pathologic flow rates. **(D,G)** Streamline angle values projected onto 3-month **(D)** and 12-month **(G)** models at physiologic and pathologic flow rates. **(E,H)** Proportion of ROIs that fall into designated angle ranges in 3-month **(E)** and 12-month **(H)** models across all flow rates.

**Figure S3:**
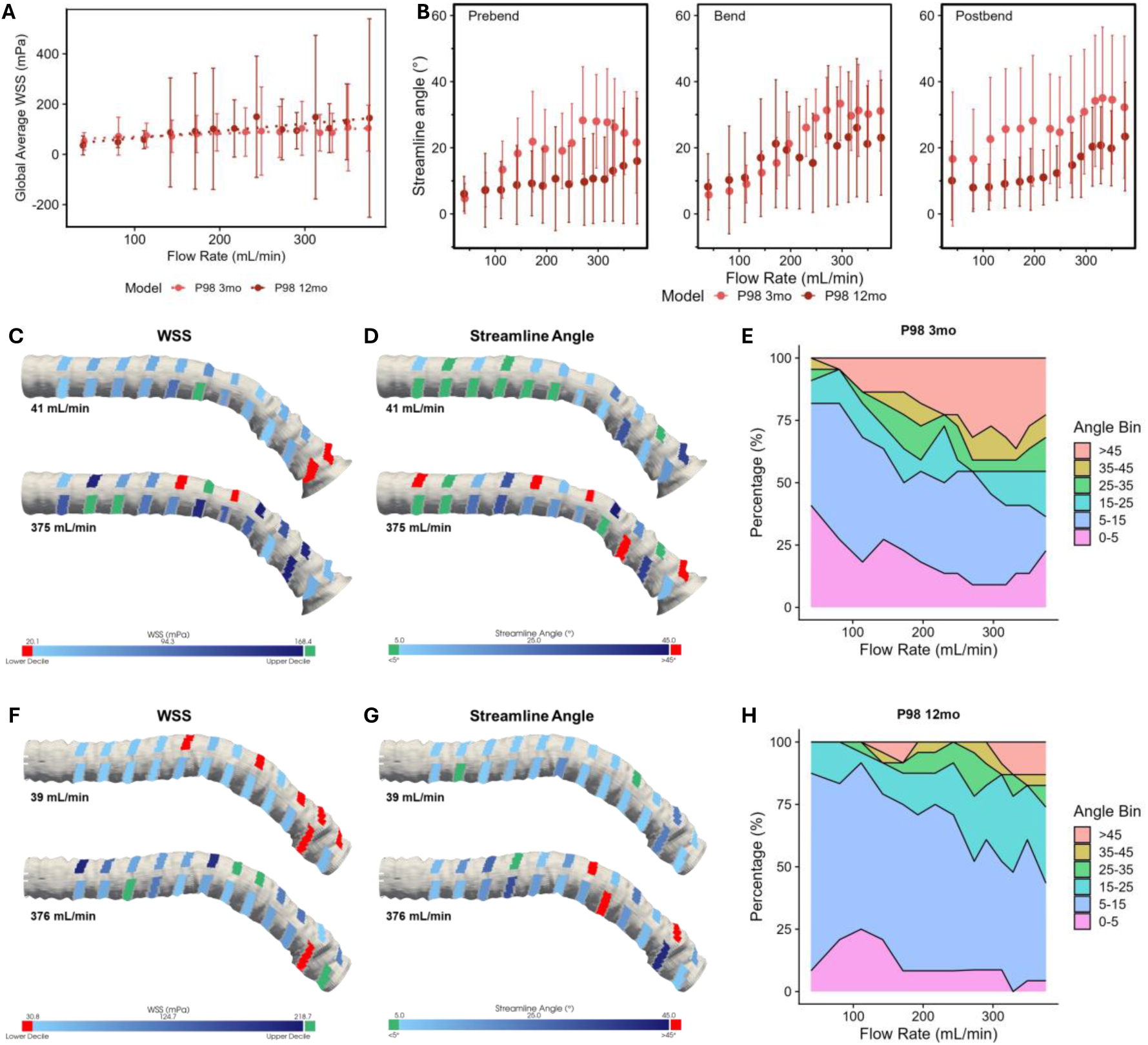
Hemodynamic profile of P98 patient models. **(A)** Global average wall shear stress (WSS) scales with flow rate. **(B)** Region-averaged streamline angle versus flow rate. Models are segmented into three regions (prebend, bend, postbend). **(C,F)** WSS values projected onto 3-month **(C)** and 12-month **(F)** models at physiologic and pathologic flow rates. **(D,G)** Streamline angle values projected onto 3-month **(D)** and 12-month **(G)** models at physiologic and pathologic flow rates. **(E,H)** Proportion of ROIs that fall into designated angle ranges in 3-month **(E)** and 12-month **(H)** models across all flow rates.

**Figure S4:**
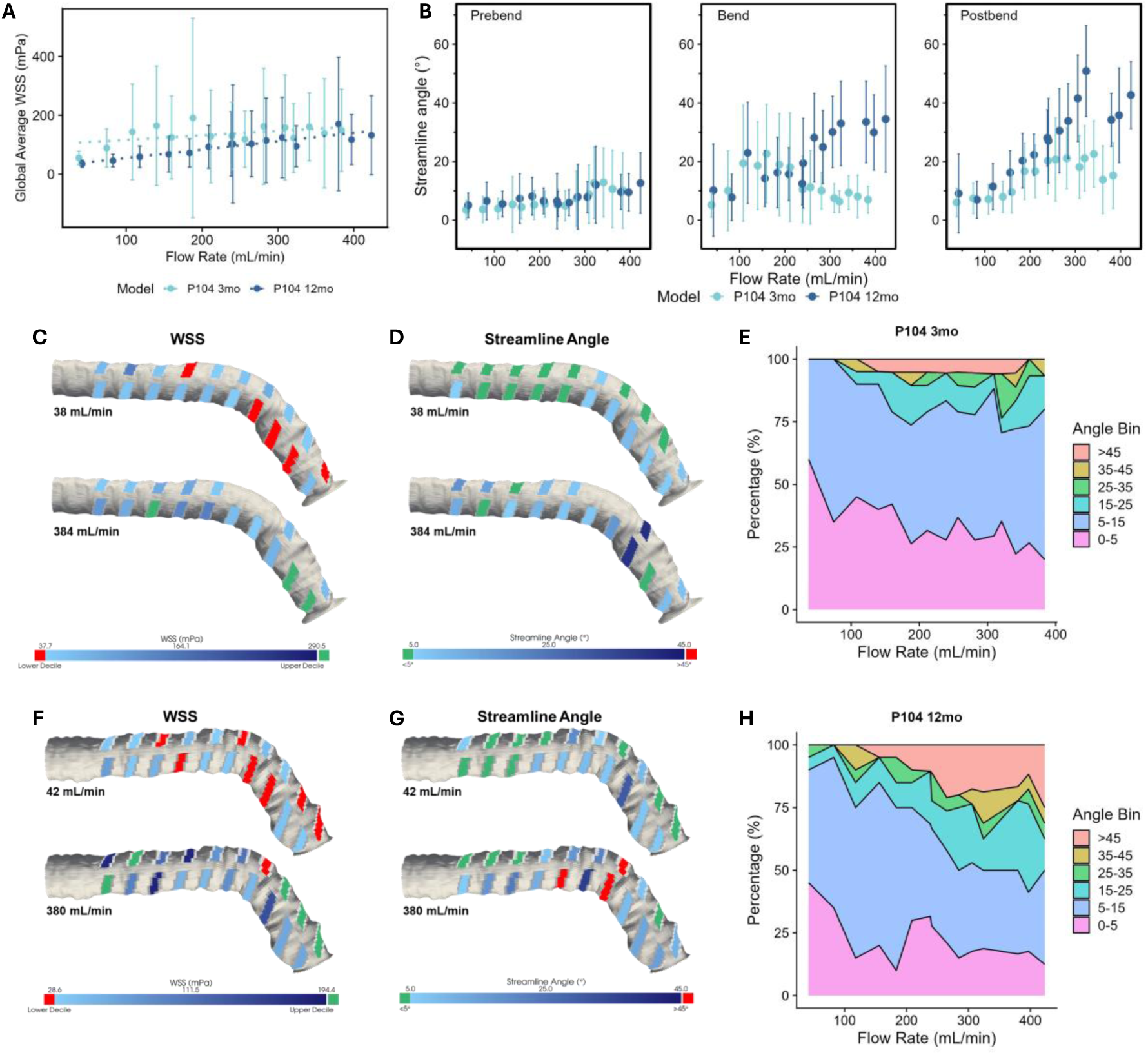
Hemodynamic profile of P104 patient models. **(A)** Global average wall shear stress (WSS) scales with flow rate. **(B)** Region-averaged streamline angle versus flow rate. Models are segmented into three regions (prebend, bend, postbend). **(C,F)** WSS values projected onto 3-month **(C)** and 12-month **(F)** models at physiologic and pathologic flow rates. **(D,G)** Streamline angle values projected onto 3-month **(D)** and 12-month **(G)** models at physiologic and pathologic flow rates. **(E,H)** Proportion of ROIs that fall into designated angle ranges in 3-month **(E)** and 12-month **(H)** models across all flow rates.

**Figure S5:**
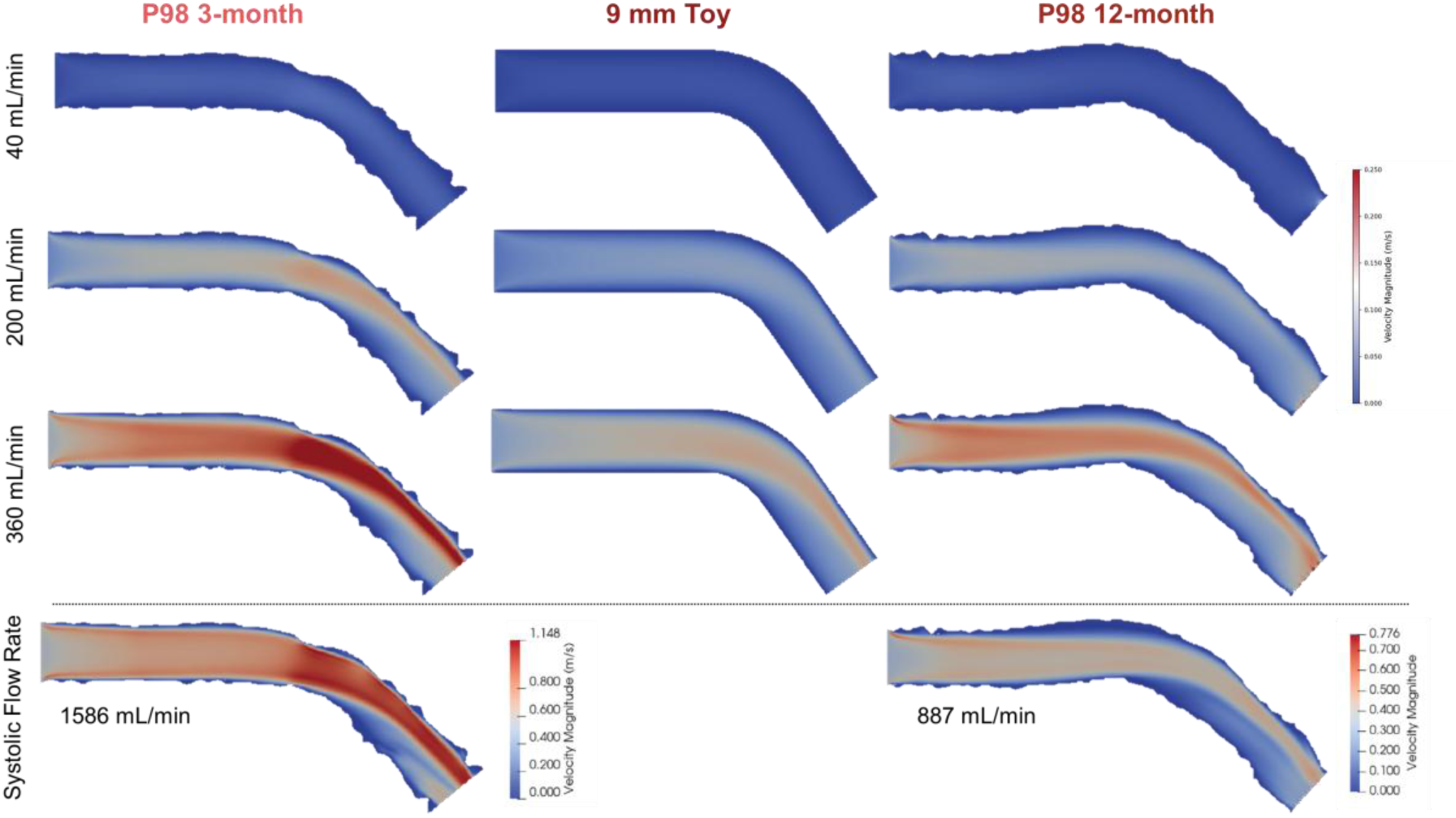
CA geometry and vessel topography affects fluid velocity field. Distribution of fluid velocity in P98 3-month, 9 mm toy, and P98 12-month models. Computational fluid dynamics simulations were performed at physiologic (40 mL/min), moderately elevated (200 mL/min), and highly elevated (360 mL/min) flow rates. For the patient models, additional simulations were performed at observed systolic flow rates.

**Figure S6:**
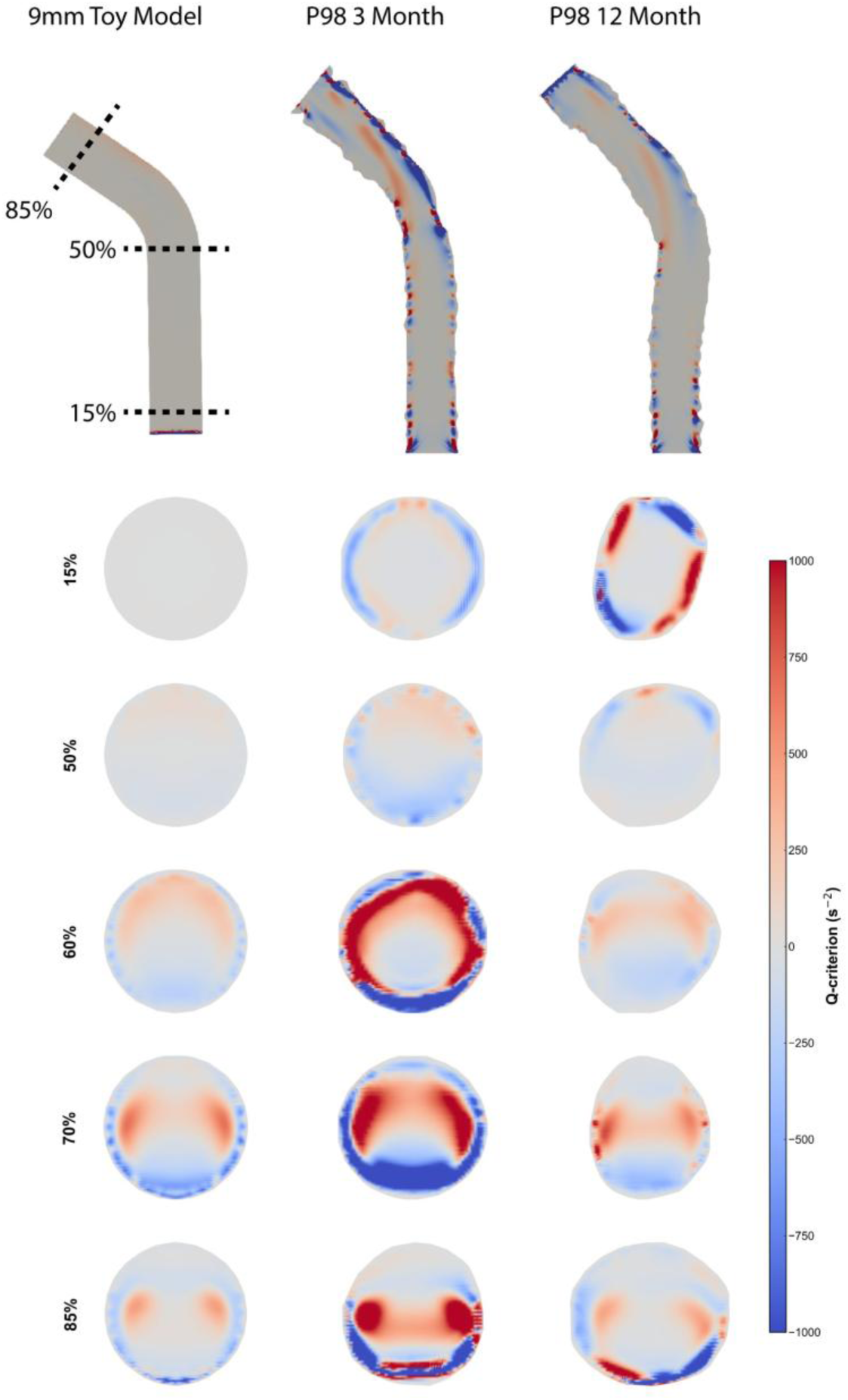
Circumferential distribution of Q-criterion in toy and patient models. Distribution of Q-criterion in P98 3-month, 9 mm toy, and P98 12-month models. Computational fluid dynamics simulations were performed at a highly elevated (360 mL/min) flow rate. Slices of the models were taken at discrete positions along the lengths of the models in the direction of flow to show the distribution of Q-criterion at relative positions.

**Movie S1: Disturbed flow develops in a quasi-aneurysm at elevated flow rates** This movie shows a video representation of data presented in **Fig. 4B**. Flow through a bulge in the vessel wall appears laminar at a physiologic flow rate (39 mL/min). At an elevated flow rate (380 mL/min), the same region harbors a recirculation eddy.

**Movie S2: Disturbed flow develops in a post-stenotic region at elevated flow rates** This movie shows a video representation of data presented in **Fig. 4D**. Flow immediately downstream of a narrowing in the vessel wall appears laminar at a physiologic flow rate (41 mL/min). At an elevated flow rate (375 mL/min), the same region harbors a large recirculation eddy.

**Table S1.**
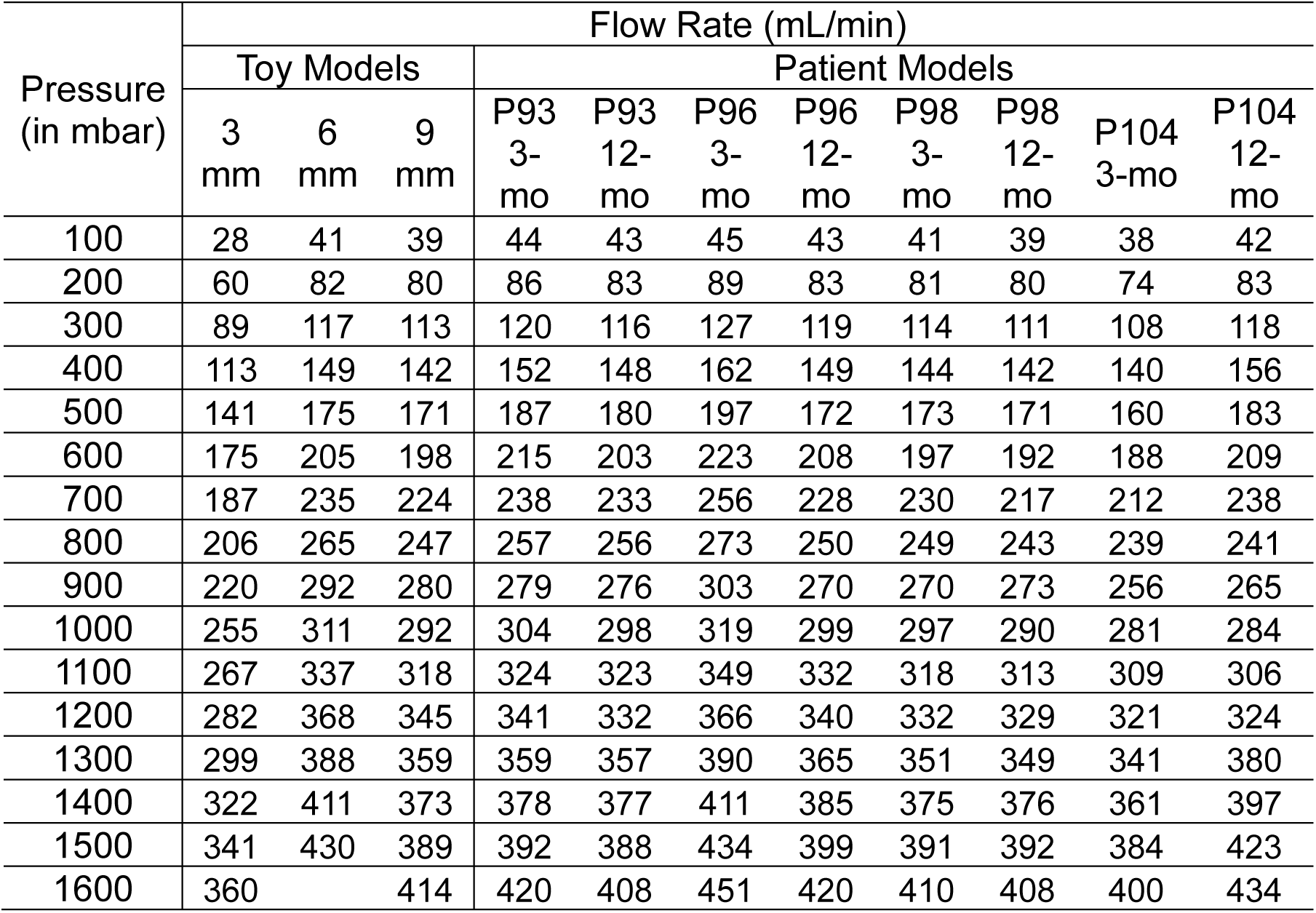
Observed flow rates for each pressure applied to inlet reservoir.

